# Predicting coral adaptation to global warming in the Indo-West-Pacific

**DOI:** 10.1101/722314

**Authors:** Mikhail V. Matz, Eric Treml, Benjamin C. Haller

## Abstract

The potential of reef-building corals to adapt to increasing sea surface temperatures is often speculated about but has rarely been comprehensively modeled on a region-wide scale. Here, we used individual-based simulations to model adaptation to warming in a coral metapopulation comprising 680 reefs and representing the whole of the Central Indo-West Pacific. We find that in the first century of warming (approximately from 50 years ago to 50 years in the future) corals adapt rapidly by redistributing pre-existing adaptive alleles among populations (“genetic rescue”). In this way, some coral populations - most notably, Vietnam, Japan, Taiwan, New Caledonia, and the southern half of the Great Barrier Reef - appear to be able to maintain their fitness even under the worst warming scenarios (at least in theory, assuming the rate of evolution is the only limitation to local coral recovery). Still, survival of the majority of reefs in the region critically depends on the warming rate, underscoring the urgent need to curb carbon emissions. Conveniently, corals’ adaptive potential was largely independent of poorly known genetic parameters and could be predicted based on a simple metric derived from the biophysical connectivity model: the proportion of recruits immigrating from warmer locations. We have confirmed that this metric correlates with actual coral cover changes throughout the region, based on published reef survey data from the 1970s to early 2000s. The new metric allows planning assisted gene flow interventions to facilitate adaptation of specific coral populations.

## INTRODUCTION

The world has been warming at an unprecedented rate for the past half century (Pachauri, Mayer, & Intergovernmental Panel on Climate Change, 2015). This brings about major ecological changes (Parmesan, 2006), among which the worldwide decline of coral reefs is one of the most alarming (O. Hoegh-Guldberg et al., 2007). Several highly cited publications have asserted that natural evolution is too slow to allow corals to adapt to global warming (O. Hoegh-Guldberg, 1999; O. Hoegh-Guldberg et al., 2007; Ove Hoegh-Guldberg, Poloczanska, Skirving, & Dove, 2017); however, this view is debatable. The scenario envisioned in these papers is “evolutionary rescue”: adaptation via the origin and spread of entirely novel adaptive mutations (Orr, 2005). This would indeed be slow, as well as poorly predictable without knowledge of such elusive parameters as mutation rate, number of potentially adaptive loci, and mutational effect size. However, the first-order evolutionary response in natural populations rarely involves new mutations; instead, it is based on adaptive alleles pre-existing in a population, collectively called “standing genetic variation” (Barrett & Schluter, 2008; Hermisson & Pennings, 2005; Savolainen, Lascoux, & Merilä, 2013). This mode of adaptation can be very rapid (Campbell-Staton et al., 2017; Lescak et al., 2015). In a metapopulation in which a species is distributed across multiple locally adapted sub-populations, there is also a possibility of rapid adaptation through “genetic rescue” (Whiteley, Fitzpatrick, Funk, & Tallmon, 2015), which involves redistribution of pre-existing locally adaptive alleles between populations through migration when conditions start to change. Several recent papers argued that this particular mode of adaptation is likely to be of major importance for reef-building corals as they adapt to warming (Bay, Rose, Logan, & Palumbi, 2017; Kleypas et al., 2016; Matz, Treml, Aglyamova, & Bay, 2018). Indeed, all coral species exist across a considerable gradient of temperatures while genetically adapting to local thermal conditions (Bay & Palumbi, 2014; Dixon et al., 2015; Palumbi, Barshis, Traylor-Knowles, & Bay, 2014) and exchanging migrants over very long distances (Ayre & Hughes, 2004; I. B. Baums, Miller, & Hellberg, 2005; Matz et al., 2018; Romero-Torres, Treml, Acosta, & Paz-García, 2018), which appears to set the perfect stage for genetic rescue (Matz et al., 2018; Torda et al., 2017).

Here, we aimed to identify factors affecting the corals’ potential to survive under warming across the central Indo-West Pacific, the ocean region where the majority of the world’s coral reefs are found (Fig. 1 A). We have extended our earlier individual-based model of metapopulation adaptation (Matz et al., 2018) to capitalize on the non-Wright–Fisher framework available in the simulation software SLiM 3.3 (Haller & Messer, 2019). This framework allows for overlapping generations, age structure within populations, age-specific mortality, and involves natural dependence of population size on fitness. The new model (https://github.com/z0on/coral_triangle_SLiM_model) was parameterized with the number of populations, reef habitat sizes, and connectivity pattern expected for a reef-building coral of genus *Acropora* (Treml, Roberts, Halpin, Possingham, & Riginos, 2015; Treml et al., 2012). Our model included a period of long-term adaptation to temperatures fluctuating around a location-specific mean followed by the onset of location-specific warming as predicted under RCP 4.5 or RCP 8.5 (Fig. S1 (Alvich, n.d.)). We have explored the influence of warming rate, population turnover rate (modulated through juvenile mortality), and genetic parameters affecting the efficiency of selection (heritability, plasticity) or amount of genetic variation (number of quantitative trait loci, mutation rate, and mutation effect size). Lastly, we have compared our results to the actual coral cover changes observed throughout the region by early 2000s (Bruno & Selig, 2007).

**Figure 1.**
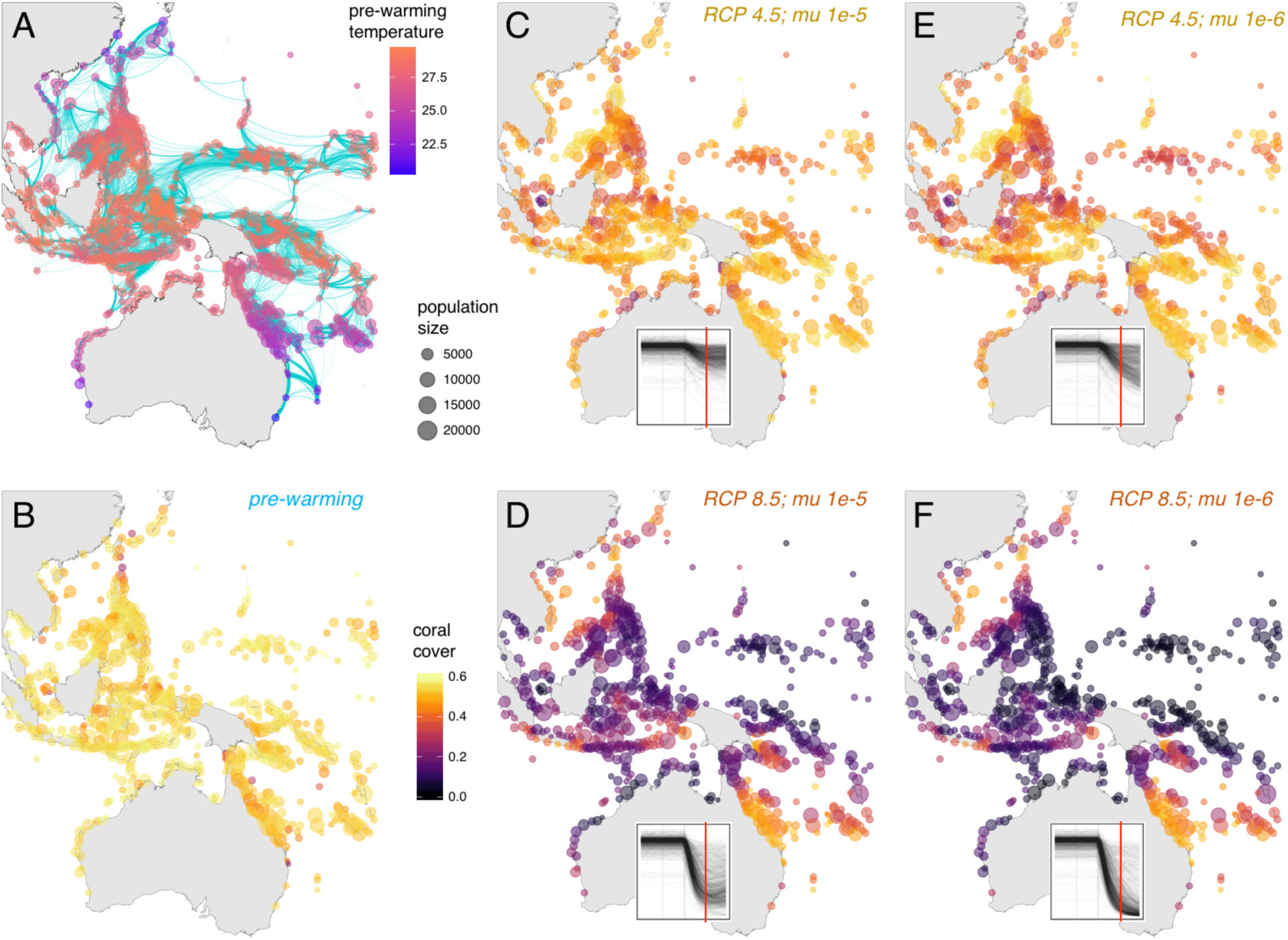
Response of coral cover (% of occupied habitat) to the first 100 years of warming depends predominantly on the warming rate and not on genetic parameters. Sizes of points on all panels indicate habitat size at each location. A: Pre-warming temperatures and migration patterns. Migration (cyan arcs) are to be read clockwise to infer direction; arc line widths correspond to 0.1-1%, 1-10%, and 10-100% probability of migration. B: pre-warming coral cover. C,D: coral cover after 100 years of warming under RCP 4.5 (C) and RCP 8.5 (D) and mutation rate 1e-5 per locus per genome replication. E,F: the same as C and D but with tenfold lower mutation rate. Insets on panels C–F show coral cover tracks for all reefs from 200 years before warming to 200 years after the onset of warming; red line marks 100 years of warming.

**Figure 2.**
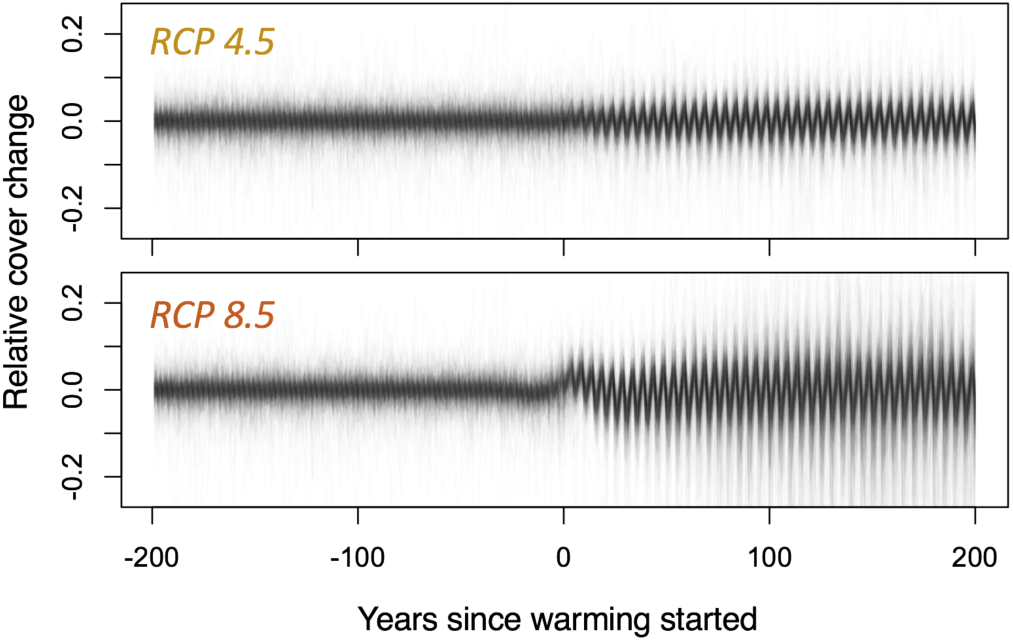
Coral cover changes (i.e, mortality-recovery cycles) in response to sinusoidal temperature fluctuations intensify during warming. Lines for all 680 populations are overlaid; each line shows deviation of the coral cover value relative to the smoothed mean.

**Figure S1.**
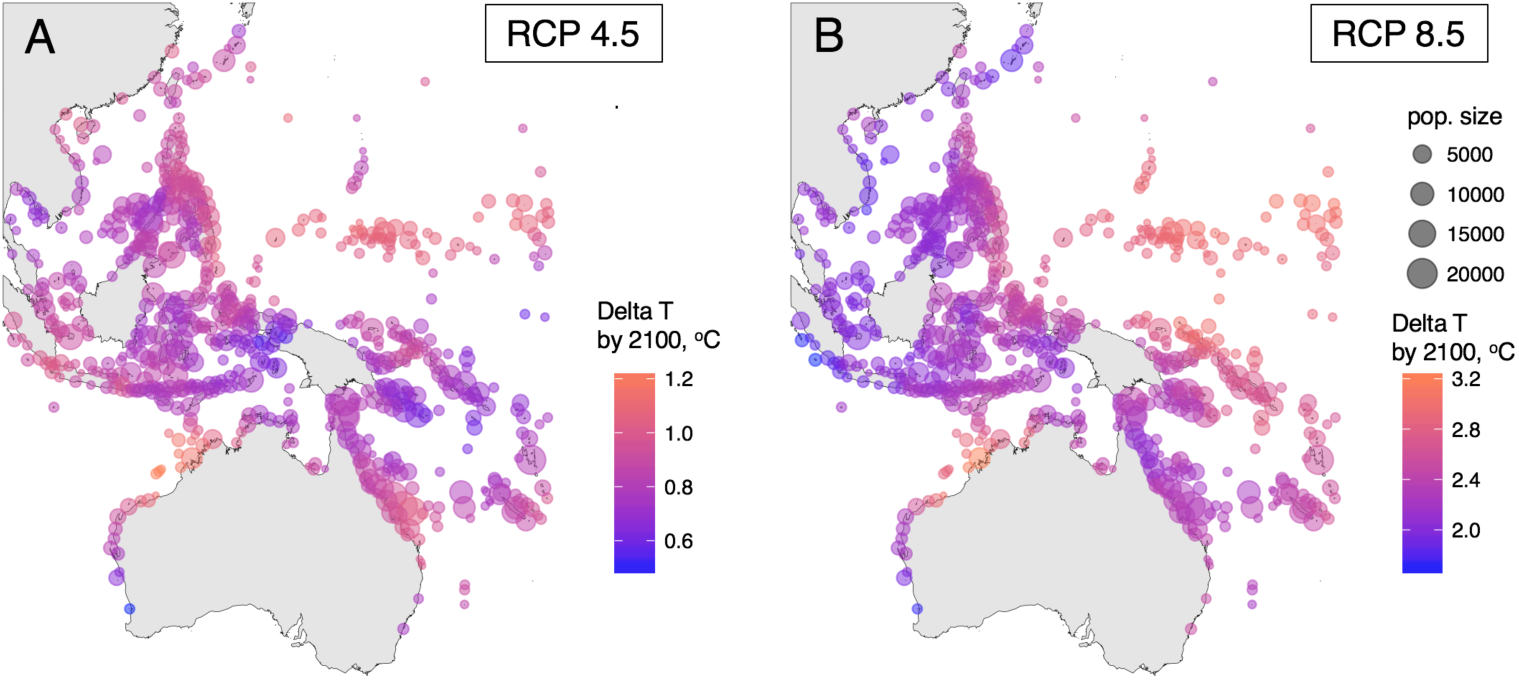
Modeled warming scenarios. Size of the points indicates carrying capacity of the population (see legend on panel B).

## MATERIALS AND METHODS

### Non-Wright–Fisher model

Compared to our earlier model (Matz et al., 2018), the new model uses a “forward migration” matrix, giving probabilities for an offspring produced at a given location to settle at all locations (including the natal one), instead of the original “immigration” matrix (giving, per location, proportions of immigrants from all locations). The non-Wright–Fisher formulation also made it possible to track the statistic that is meaningful for coral ecology – coral cover – as a proportion of the available carrying capacity occupied. The model also outputs (per population) a set of metrics: mean phenotype, mean fitness, number of segregating mutations at the QTLs, standard deviation of breeding value, mean age of adults, number of adults, and adult mortality in the last generation. The model code (https://github.com/z0on/coral_triangle_SLiM_model) reads habitat sizes, the migration matrix, and environmental settings from external files, while the major population-genetic parameters (see next paragraph) are supplied as external arguments. This makes the model easy to repurpose for any metapopulation-evolution scenario.

### Main parameter settings

Other than transitioning to the non-Wright–Fisher framework, the model remained very similar to the original one (Matz et al., 2018) in structure. We used conservative parameters both in terms of allowed genetic variation and factors affecting the efficiency of selection. We assumed 100 additive quantitative trait loci (QTLs) affecting an individual’s thermal optimum, with new mutations having subtle effects following the distribution *N* (0, 0.03°C). With these settings and in the absence of selection the mutational process would have resulted in standard deviation in the thermal optimum of only 0.1°C in a population. The genetically determined thermal optimum (breeding value) of an individual was calculated as the sum of the effects of all QTLs, and the actual phenotype was then computed by adding a random value drawn from *N*(0, 0.5°C) to model imperfect heritability. The fitness of an individual was computed based on the difference between its phenotype and the environmental temperature at its location, given a Gaussian fitness curve with a standard deviation of 1; this fitness function implies that an individual’s fitness would drop by about 40% if its phenotype were mismatched with the local temperature by 1°C.

### Reproduction and age-related mortality

Each *Acropora* individual produces 10^5^–10^6^ eggs per yearly spawning event but the vast majority of these offspring die as larvae or very young recruits without giving rise to a new colony. We assumed that each adult coral produces just a single surviving recruit per year, and that recruit still has a 10-fold higher chance of mortality in its first year compared to an adult of the same phenotype. Second-year juveniles had a 1.5-fold higher chance of mortality than adults; in the third year the surviving corals become reproducing adults (Baria, dela Cruz, Villanueva, & Guest, 2012). This age-specific mortality profile resulted in average adult coral age, prior to warming, of approximately 10 years, which is reasonable for fast-growing acroporids.

### Habitat size and genetic variation

The critical parameter determining the total amount of genetic variation available to selection is the product of population size and mutation rate. Here, we assumed that the smallest reefs (completely enclosed within a single 10×10km cell) could contain 100 corals, and larger reefs had more carrying capacity, proportionally to the number of 10×10km cells they occupied (up to 20,000 corals per reef). While these numbers are on par with genetic estimates of effective population sizes (Matz et al., 2018), they are much lower than census sizes. We kept our population sizes low to keep the model conservative (i.e., limiting for adaptation) and also faster-running, but compensated for it by assuming a mutation rate on the upper boundary of values reported in the literature, 1e-5 per QTL per generation (Barton & Turelli, 1989). In our earlier model (Matz et al., 2018), this mutation rate allowed for indefinite adaptation based on novel adaptive mutations (evolutionary rescue).

### Environment

Location-specific temperatures were based on the mean yearly temperature at each location (Fig. 1A) during the genetic equilibration period (see below), after which warming was imposed with a location-specific rate predicted under either RCP 4.5 or RCP 8.5 (Fig. S1 (Alvich, n.d.)). We explored three models of temperature variation (to which warming was added): constant, fluctuating as a sine wave with a period of 5 years and amplitude of 0.5°C (to approximate El Niño cycles (Quinn, Taylor, & Crowley, 1993)), and random temporally uncorrelated fluctuations with an amplitude drawn from a normal distribution using the standard deviation of temperatures empirically observed at each location. These three environmental models yielded nearly identical results, and so all the figures presented here correspond to the sinusoidal model.

### Parameter variations

For each parameter we tried a different setting in addition to the setting in the “main run”, summarized in Fig. 3. The “Fewer QTLs” scenario involved 10 QTLs (instead of 100) with 3-fold higher possible mutational effects; i.e., drawn from *N* (0, 0.1°C) instead of *N* (0, 0.03°C). This adjustment of the mutation effect distribution was done to preserve the mutational genetic variation, which is the square of the standard deviation of mutation effects times the number of QTLs. In the “Low juvenile mortality” scenario, first-year recruits were only two times (as opposed to ten times) more likely to die as adults, and in the second year their survival was only 10% lower than adults. This scenario resulted in an average adult age of 6.5 years. The “Lower plasticity” scenario used a narrower fitness function, implying that the fitness of an individual would drop by 86% (instead of 40%) when its phenotype mismatches the environment by 1°C. The “Higher heritability” scenario had a smaller random value added to the breeding value when computing phenotype; this resulted in a heritability of 0.4 in locally adapted populations prior to warming, compared to 0.15 in the main run. The “Lower mutation rate” scenario used a tenfold lower rate, and the “Lower mutation effect” scenario drew effect sizes from *N* (0, 0.01°C), threefold lower than in the main run. Populations in the “Larger populations” scenario were twofold larger, with two corals per km^2^.

**Figure 3.**
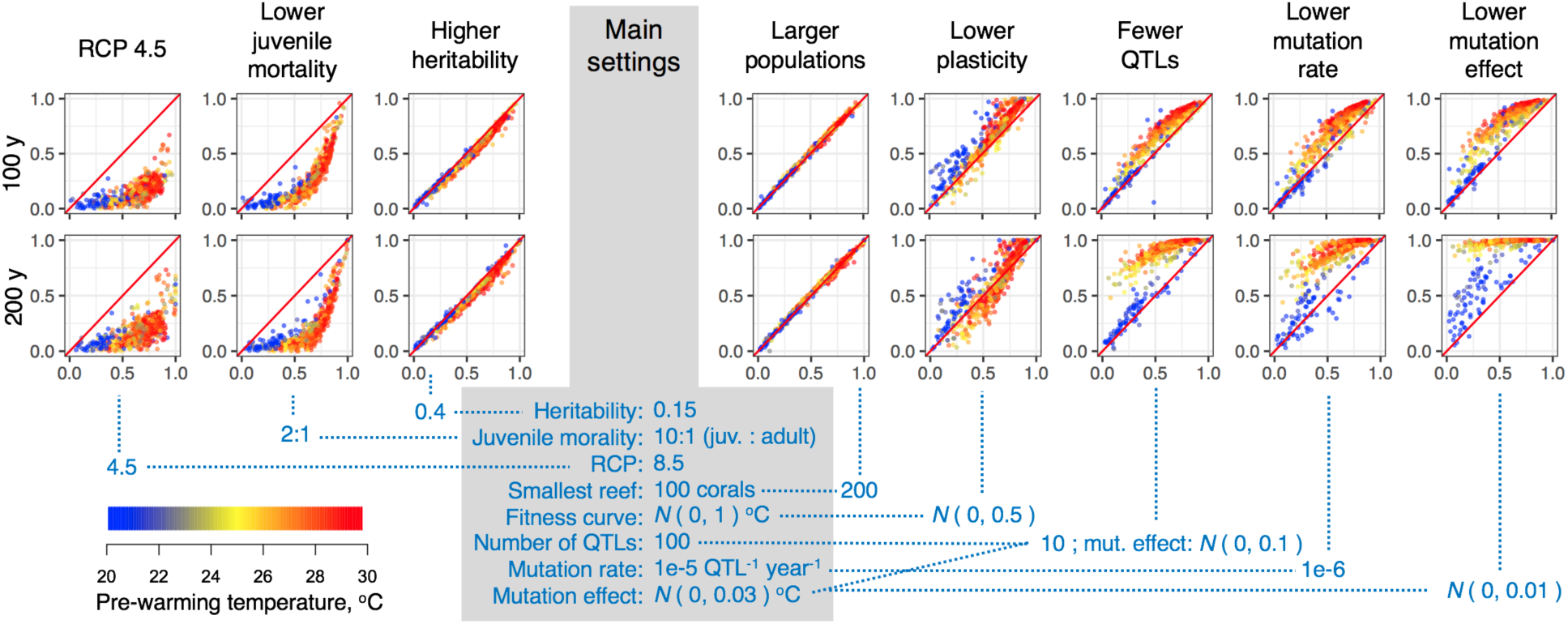
Comparisons of coral cover declines under different parameter settings, after 100 (first row of plots) and 200 (second row of plots) years of warming. Coral cover declines under alternative parameter settings (y axis) range from 0 (no decline) to 1 (extinction), and are plotted against declines observed under the “main settings” model (x-axis). Each model is different from the main in a single parameter indicated under the plots except the “Fewer QTLs” scenario, which had fewer QTLs and elevated mutation effect to keep the amount of mutational genetic variation constant. Colors indicate mean pre-warming temperature of the reef (see legend). The red line on each panel is 1:1 correspondence. Declines in the first 100 years (first row of plots) are similar across parameter settings, with the notable exceptions of lower warming rate (RCP4.5) and lower juvenile mortality, both of which lead to faster population turnover relative to the rate of warming. After 200 years (second row of plots), many mid- and high-temperature reefs decline dramatically under settings that limit the influence of new adaptive mutations (“Fewer QTLs”, “Lower mutation rate”, “Lower mutation effect”), preventing recovery via evolutionary rescue.

### Migration

The movement of individuals among sub-populations of *Acropora* was estimated using our biophysical model of larval dispersal for the Indo-Pacific (Treml et al., 2015, 2012). This model incorporates hydrodynamic data (1/12° & daily resolution for 1992-2012 (“HYCOM Data Acknowledgement Statement,” n.d.)), coral reef habitat maps (“Ocean Data Viewer,” n.d.), and biological parameters for coral larvae. In this model, dispersing larvae were represented by their i) spawning time following two summer full moons, ii) larval settlement window from 6 days to 60 days (maximum larval duration (Connolly & Baird, 2010)), iii) daily larval mortality (5% (Connolly & Baird, 2010)) and iv) passive behavior while distributed in the top 10 meters of the ocean. In this model, larvae are moved throughout the seascape following spawning using an efficient and 4^th^ order accurate advective transport scheme (Smolarkiewicz, 1983). A detailed model description and sensitivity analysis is available in (Treml et al., 2012). The results from the dispersal simulations were used to create a long-term average forward transition matrix quantifying the likelihood that larvae spawned at a source reef survive and settle to all potential reef sites (including the natal source patch).

### Genetic equilibration

The model used the same stepwise procedure to rapidly achieve genetic equilibrium as the original model used (Matz et al., 2018). The first 2000 years were run with population sizes 25-fold smaller but mutation rate 25-fold higher than target values, followed by 2000 years of 10-fold smaller population sizes / 10-fold higher mutation rate. The remaining years were run at the target population sizes and mutation rates; the warming began at year 5500. In this way, the product of population size and mutation rate that governs the overall genetic variation is constant throughout the simulation, but the genetic equilibrium is approached substantially faster due to the smaller population size at the beginning of the simulation. We have confirmed that the genetic variation stays constant in the last 200 years preceding warming (Fig. S2), so the adaptation to warming starts from a state of genetic equilibrium.

### Model runs and results processing

The analysis focused on the 400-year window centered on the start of warming. All models were run five times with different random seeds; the decadal averages for each run were averaged among runs. The coral cover per site was computed as the number of adults divided by the carrying capacity of the site. The coral cover response (Fig. 3) was computed as the difference between coral cover after 100 or 200 years of warming and coral cover in the pre-warming decade, divided by the pre-warming cover. Reef declines plotted in Fig. 2 are absolute values of responses for reefs that experienced declines (more than 95% of all reefs).

### Matching with actual reef survey data

Bruno and Selig (Bruno & Selig, 2007) have compiled coral surveys across the Indo-Pacific from the 1970s to early 2000s, to quantify region-wide coral declines. To match the locations of surveyed and simulated reefs, we clustered all reefs into groups within distance corresponding to one degree of latitude (111 km) from each other, and selected clusters that (i) contained both simulated and real reefs, (ii) contained real reefs surveyed >15 years apart, and (iii) contained a reef survey completed after 2000. All the real reef data within each cluster were then used to compute a regression coefficient of coral cover against year. Essentially, this analysis treats all surveys of reefs within a local cluster as replicate surveys of the same reef (very few individual reefs were actually surveyed repeatedly over a long period). These “real-change” regression coefficients were compared to the mean environmental predictor value, or to the mean model-predicted coral cover response, across virtual reefs within the same cluster.

## RESULTS

Prior to warming, adult coral cover at each site stabilized at similar levels across most models (Fig. S2), indicating that populations were nearly equally successful at local adaptation irrespective of most parameter settings. There were a few exceptions. The cover was lower under low plasticity, which is explained by lower mean population fitness under this setting, and under low juvenile mortality, which leads to a smaller proportion of adults in each population. Conversely, the cover was higher under higher heritability due to the higher mean population fitness attainable under this setting. The within-population standard deviation of breeding value (square root of genetic variation) stabilized at approximately 0.2°C (Fig. S2). The stability of this value in pre-warming generations indicates that genetic equilibrium has been reached. Notably, pre-warming genetic variation was similar irrespective of the model’s settings (Fig. S2), indicating that the same amount of standing genetic variation was available to selection at the onset of warming. The stability of this value irrespective of settings that affect the number of new mutations arising each generation (population size or mutation rate) suggests that standing genetic variation in this system predominantly depends on migration-selection balance rather than on mutational input.

**Figure S2.**
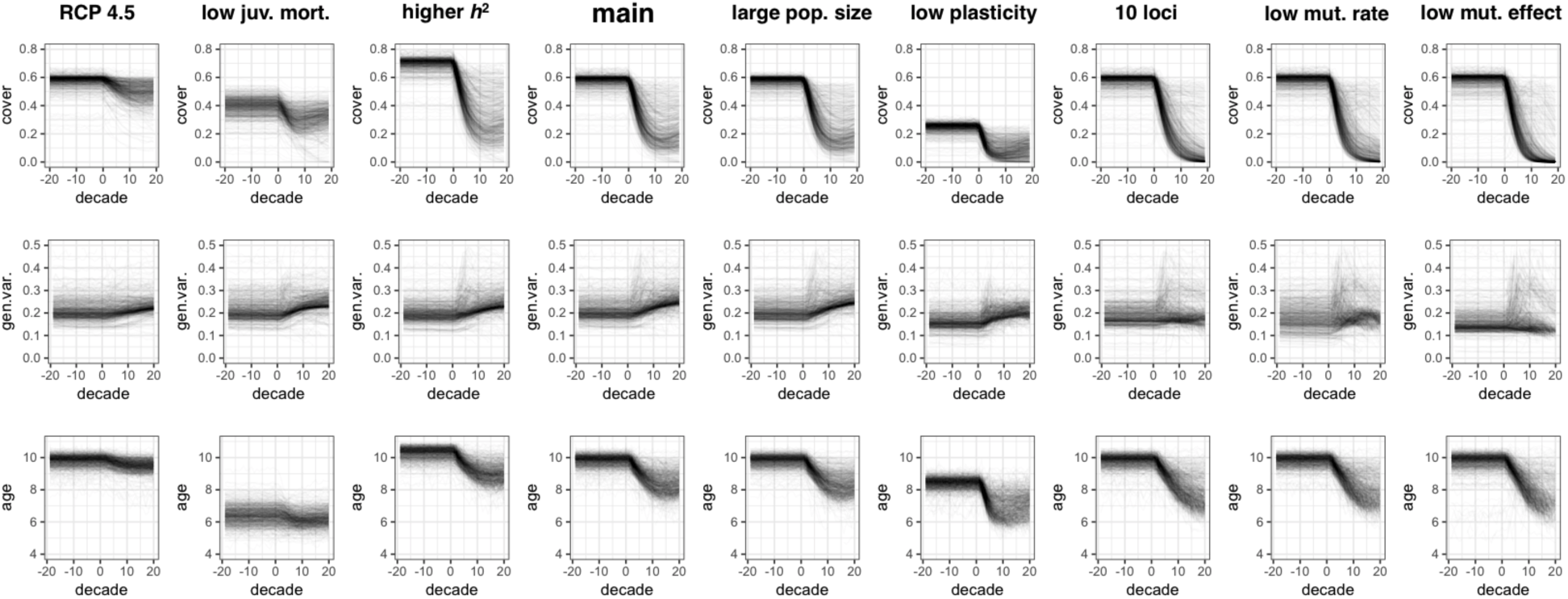
Profiles of coral cover (% of occupied carrying capacity, top row), standard deviation of breeding value (i.e., square root of genetic variation, middle row), and mean adult age (bottom row) under different parameter settings. The settings are identified above the columns and correspond to settings listed in Fig. 3. The *x*-axis is decade since the start of warming; the *y*-axis is the value at a particular reef. On each panel, traces for all 680 reefs are overlaid. Note that in all cases the most dramatic change happens within the first 10 decades of warming.

The mean age of adult corals was also similar – about 10 years – both among populations and model runs (Fig. S2), which is a reasonable number for fast-growing *Acropora* corals (Baria et al., 2012). There were three exceptions: the mean age was higher under high heritability, lower under low plasticity, and much lower under low juvenile mortality. Higher heritability leads to older corals because of their higher average fitness and therefore lower yearly risk of mortality, and conversely, younger average age under low plasticity is explained by lower fitness and higher risk of mortality. Younger average age under low recruit mortality is a direct consequence of more young corals surviving in the first year.

The spatial pattern of reef responses to warming was qualitatively similar across all model settings, varying mostly in intensity (Fig. 1, Fig. 3, <u>Supplemental video 1</u>, https://www.dropbox.com/s/ffv2zb164gwosld/slim_triangle_v4.mp4) By far the most dramatic changes happened in the first century of warming (Fig. 1 C–F, insets; Fig. 3). Supporting the observation from our previous model (Matz et al., 2018), warming resulted in an increase in magnitude of coral cover response to thermal fluctuations, i.e., more severe coral mortality episodes in response to heat waves (Fig. 2; this can also be seen in <u>Supplemental video 1</u>, https://www.dropbox.com/s/ffv2zb164gwosld/slim_triangle_v4.mp4, as intensifying pulses of darker color).

Corals did much better overall under the slower warming rate (RCP 4.5, Fig. 1 C, E) and low juvenile mortality (Fig. 3). With all other parameter settings the responses were generally similar in the first 100 years (Fig. 3), which is expected since the initial adaptive response is based predominantly on standing genetic variation (i.e., it is mostly genetic rescue – redistribution of pre-existing adaptive alleles among populations), and this variation stabilizes at similar levels across parameter settings (Fig. S2). Only when standing genetic variation starts running out do other settings, especially those affecting new mutations – fewer QTLs, lower mutation rate, and lower mutation effect size – begin to have substantial influence (Fig. 3). These settings diminish the chance of adaptation based on novel mutations (evolutionary rescue) and prevent coral cover from stabilizing and recovering, especially at mid- and high-temperature reefs (Fig. 3).

Encouragingly, decline was not the universal response among reefs. Consistently across all parameter settings, reefs located in the Northwest and Southeast parts of the modeled region were largely unaffected, even under rapid warming and low mutation rate setting (Fig. 1). The strongest predictor of coral cover response to warming was pr05: the proportion of recruits that come from locations that are at least 0.5°C warmer (Fig. 4). This parameter essentially quantifies the potential of a reef to undergo genetic rescue via immigration of warm-adapted alleles. It alone explains 49% of variation in reef response under RCP 4.5 and 68% of variation under RCP 8.5 (Fig. 4 B, C). For comparison, the next-best predictor – mean reef temperature under RCP 8.5, Fig. 3 – explains only 4% of variation in addition to pr05, or 42% of variation on its own (it is highly correlated with pr05).

**Figure 4.**
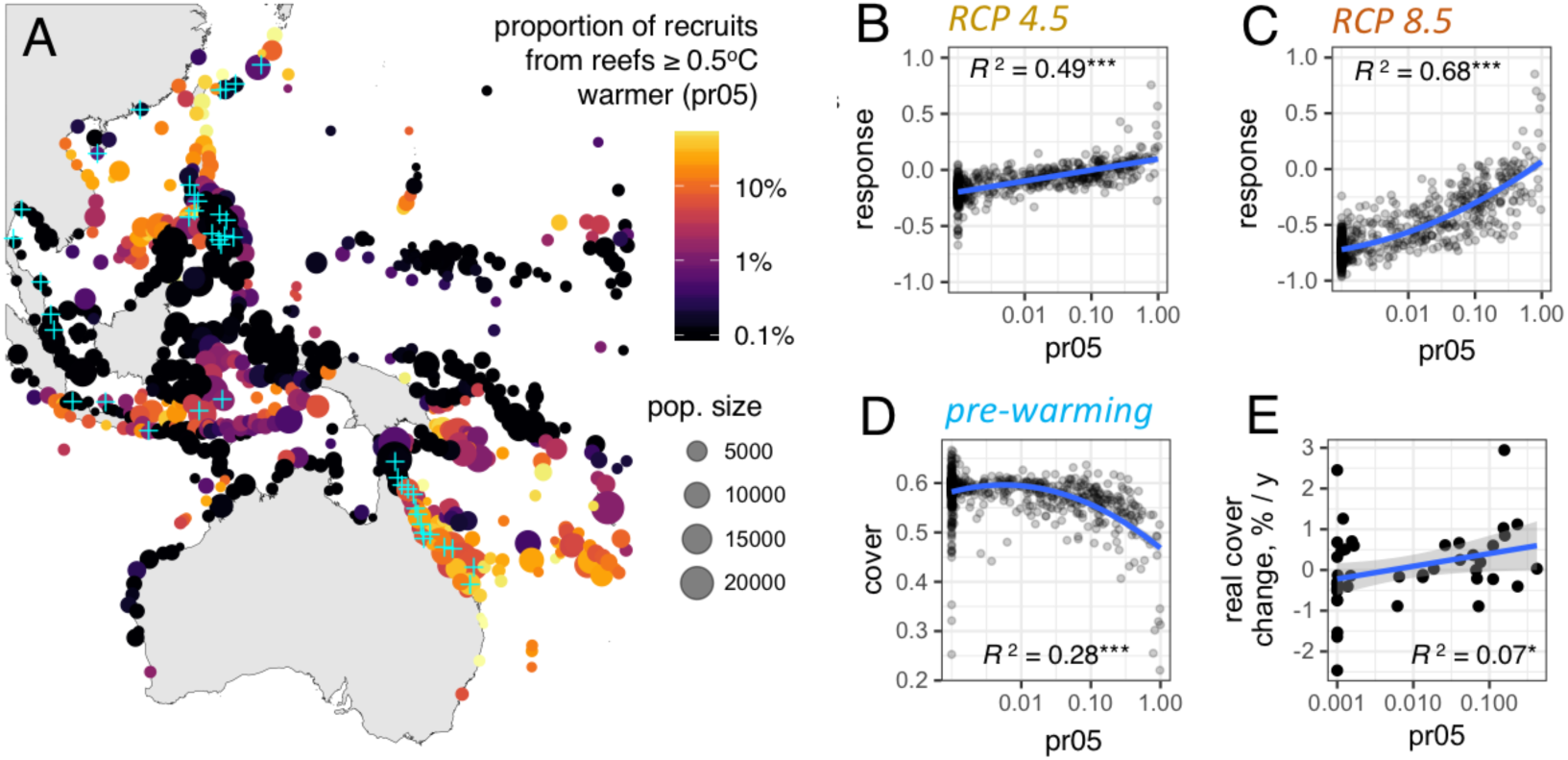
Immigration from warmer locations predicts reef response to warming. A: Map of pr05, the proportion of recruits immigrating from reefs that are at least 0.5°C warmer. Note the similarity of this pattern to the pattern of coral cover after 100 years of rapid warming (Fig. 1 D, F). The locations for which the actual long-term coral cover data were available (Bruno & Selig, 2007) are marked by cyan crosses. B, C: relationship between pr05 and coral cover response after 100 years of warming (negative = decline, positive = increase), under RCP 4.5 (B) and RCP 8.5 (C). D: relationship between pr05 and pre-warming coral cover: too much immigration from warmer locations interferes with local adaptation, but this effect is mild unless immigration is very high. E: correlation of pr05 and actual coral cover change over 15 years across the turn of the century, based on data from (Bruno & Selig, 2007). On panels B-E, blue lines are regressions against log pr05 (linear on B and E and polynomial with degree 2 on C and D). 95% confidence interval on panel E is grey, on B-D it is contained within the width of the line. *** *p* < 1e-10; * *p* = 0.04.

The influx of immigrants from warmer reefs is expected to interfere with local adaptation during the pre-warming period (Ronce & Kirkpatrick, 2001), and indeed, we see a negative relationship between pr05 and pre-warming coral cover (Fig. 4 D). Still, pr05 on the order of 10% leads to relatively minor maladaptation pre-warming while fully protecting a reef from decline under RCP4.5 (Fig. 4 B) and alleviating half of the decline under RCP 8.5 (Fig. 4 C).

**Figure S3.**
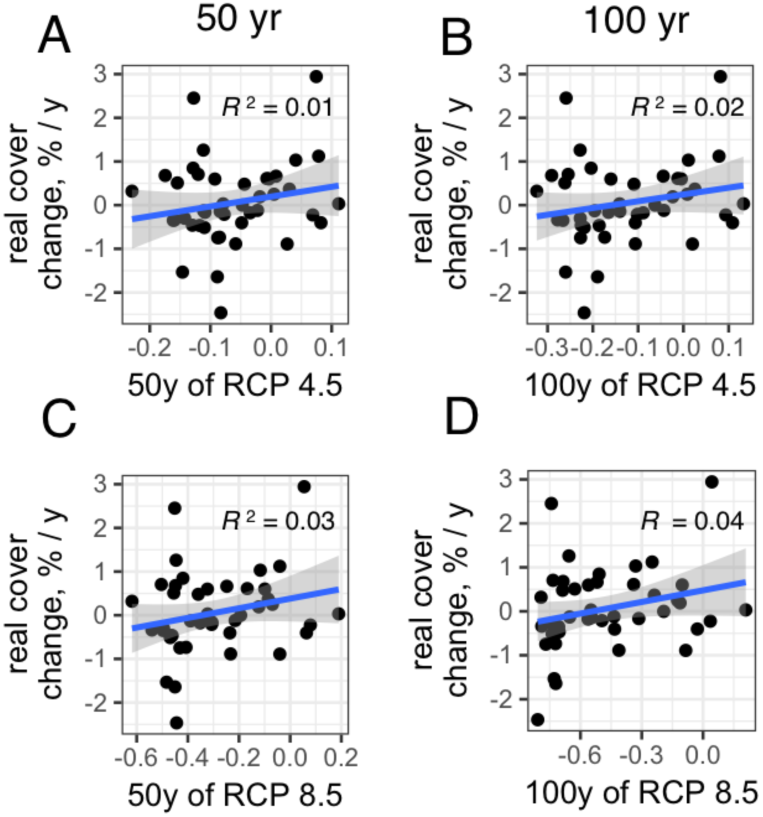
Relationship between model-predicted coral cover responses and actual coral cover change over 15 years across the turn of the century (Bruno & Selig, 2007). None of the correlations is statistically significant, although the model predictions under RCP 8.5 scenario (C, D) are slightly more similar to the actual changes.

Finally, pr05 was found to significantly correlate with the actual coral cover changes observed throughout the region (Bruno & Selig, 2007) (Fig. 4 E). At the same time, the correlation between real changes and model-predicted changes either at 50 or 100 years of simulated warming was not significant (Fig. S3). This is not very surprising since the model is an abstraction of reality, not accounting for many relevant factors (most notably, ecological interactions, as we discuss below). Yet, it appears that our model did help to identify pr05 as an important environmental predictor of coral resilience.

## DISCUSSION

The world is already about 50 years into the warming scenario that we model (Pachauri et al., 2015). Can evolution rescue corals from global warming? Yes and no. As long as we are willing to assume that coral populations are genetically adapted to their local thermal regimes (Bay & Palumbi, 2014; Dixon et al., 2015; Palumbi et al., 2014), our model indicates that evolution is going to protect at least some reefs from extinction for quite a while. Most notable of those are reefs at the latitudinal range edge: Mid- and Southern Great Barrier Reef, New Caledonia, Vietnam, Taiwan, and Japan (Fig. 1 D, F). Encouragingly, this protective effect was observed even under rapid warming and limiting settings for novel genetic variation (Fig. 1 F). The surprising resilience of these reefs in our model is explained by the fact that they are major downstream accumulation sites for adaptive genetic variation, drawing on heat-tolerant alleles immigrating from warmer reefs throughout the whole region (Fig. 4). At least in theory, slow rate of evolution should not be a problem there.

At the same time, many other regions, especially those that are already warm and don’t receive immigrants from yet warmer places (Fig. 4), are much more prone to dramatic declines due to warming, even under mutation rates potentially allowing for indefinite evolution (Matz et al., 2018) (Fig. 1 C, D). The state of these reefs critically depends on the single most important parameter: the rate of warming relative to the rate of population turnover. Slower warming (RCP 4.5) and low juvenile mortality (which leads to faster population turnover, Fig. S2) can dramatically offset declines of these reefs. Under warming rates predicted under RCP 4.5 (Fig. S1 A), which assumes reduction of greenhouse gas emissions and stabilization of the greenhouse effect by 2100, even the worst reef declines do not exceed 30–50% (depending on other model settings) within the first two centuries of warming (Fig. 1 C,E). In contrast, under the “business-as-usual” RCP 8.5 scenario the majority of reefs near the equator and in Western Australia decline to near-extinction within the first 100 years (Fig. 1 C, E). While our model is likely conservative in terms of allowed rate of accumulation of novel genetic variation due to low assumed population size, and so these declines might be somewhat offset by new adaptive mutations (evolutionary rescue), we believe that the model correctly reconstructs the general shape of the initial response based on standing genetic variation.

It is important to emphasize that our model is purely population-genetic in the sense that it assumes that any reef can recover without impediment as long as there is immigration and recruitment. We do not account for possible ecological feedbacks that might limit reef recovery, such as change in migration rates with warming (Munday et al., 2009), competition with algae (McCook, Jompa, & Diaz-Pulido, 2001), disease (Bruno et al., 2007), or transition of some coral predator to devastating boom-and-bust population cycles, as happened with the crown-of-thorns starfish (Kayal et al., 2012). We also don’t account for the increase in storm severity (Emanuel, 2005), which is a major destructive force for many Indo-Pacific reefs (De’ath, Fabricius, Sweatman, & Puotinen, 2012). It is also important to note that for corals that mature and grow slower than acroporids modeled here adaptation would be progressively unlikely, since slower population turnover rate impairs adaptation just as much as the faster warming rate (Fig. 3). All this means that even the best-protected reefs (according to our model) are still vulnerable to climate change. Although in theory they should be able to evolve rapidly enough, they remain prone to all the other sources of mortality to which global warming contributes. Essentially, our model provides a best-case scenario that can be used as a baseline to test for the role of other factors in reef decline.

Should we consider helping corals evolve? While natural selection will always be much more efficient than any lab-based selection because it has access to the vastly broader standing genetic variation in nature, our results suggest one intervention that might help: facilitating genetic influx from warmer locations, to raise the local pr05 (Fig. 4). This type of intervention is called “assisted migration” (Haller, Mazzucco, & Dieckmann, 2013) or “assisted gene flow” (Aitken & Whitlock, 2013), and would make particular sense on reefs that don’t receive any natural immigrants from warmer locations. Using cryopreserved sperm from warmer reefs to fertilize local eggs and outplanting the juveniles would introduce otherwise inaccessible adaptive alleles into the local population (Baums et al., 2019). While earlier works have proposed similar interventions (Dixon et al., 2015; Kleypas et al., 2016; Matz et al., 2018), our current results suggest the scale on which it has to be done. While even a small increase in pr05 already lowers the risk of reef decline (Fig. 4 B, C), tangible effects are only observed when pr05 is on the order of several percent or higher. This means that one would have to outplant warm-adapted recruits in numbers approaching 5–10% of total natural recruitment, which may or may not be realistic depending on the coral species and the ocean basin. Our model can be used to estimate the efficiency of such effort on specific reefs.

All that said, by far the most helpful thing that we could do for coral reefs would be to curb greenhouse gas emissions to push the global warming trajectory closer to the RCP 4.5 scenario. According to the model presented here, slowing down of the warming rate would bring the most substantial relief to struggling corals.

## Supporting information

Movie of coral cover change during 200 years of warming, under four different scenarios corresponding to Fig. 1 C-F of the manuscript.

## ACKNOWLEDGEMENTS

We are grateful to John F. Bruno for providing the compilation of coral cover data to compare our model to real observations. This work was supported by the National Science Foundation grant OCE-1737312 to M.V.M. This project relied on the high-performance computing resources of the Texas Advanced Computing Center.

